# Identification of regulators of poly-ADP-ribose polymerase (PARP) inhibitor response through complementary CRISPR knockout and activation screens

**DOI:** 10.1101/871970

**Authors:** Kristen E. Clements, Anastasia Hale, Nathanial J. Tolman, Claudia M. Nicolae, Anchal Sharma, Tanay Thakar, Xinwen Liang, Yuka Imamura Kawasawa, Hong-Gang Wang, Subhajyoti De, George-Lucian Moldovan

## Abstract

Inhibitors of poly-ADP-ribose polymerase 1 (PARPi) are highly effective in killing cells deficient in the homologous recombination (HR) DNA repair pathway, such as those lacking BRCA2. In light of this, PARPi have been utilized in recent years to treat BRCA2-mutant tumors, with many patients deriving impressive clinical benefit. However, positive response to PARPi is not universal, even among patients with HR-deficient tumors. Here, we present the results of three genome-wide CRISPR knockout and activation screens which provide an unbiased look at genetic determinants of PARPi response in wildtype or BRCA2-knockout cells. Strikingly, we reveal that depletion of the histone acetyltransferase TIP60, a top hit from our screens, robustly reverses the PARPi sensitivity caused by BRCA2 deficiency. Mechanistically, we show that TIP60 depletion rewires double strand break repair in BRCA2-deficient cells by promoting 53BP1 binding to double strand breaks to suppress end resection. Our work provides a comprehensive set of putative biomarkers that serve to better understand and predict PARPi response, and identifies a novel pathway of PARPi resistance in BRCA2-deficient cells.

## Introduction

Identifying vulnerabilities specific to cancer cells remains the central goal of much research in the field of cancer biology; when successful, this line of research can significantly improve patient survival. In 2005, two groups discovered that the very mutations that contribute to tumorigenesis in some patients—mutations in the homologous recombination (HR) proteins BRCA1 and BRCA2—also sensitized cells to treatment with inhibitors of poly-ADP-ribose polymerase 1 (PARPi).^1,2^ The classical model explaining this synergy proposes that these drugs induce double-strand DNA breaks which cannot be repaired in BRCA-deficient cells.^1,2^ In the following years, PARPi have shown remarkable promise; indeed, the FDA has approved PARPi for use in the clinic based on multiple clinical trials demonstrating an impressive improvement in progression free survival (PFS).^3,4^ For example in one study, olaparib treatment in ovarian cancer patients harboring a germline BRCA1/2 mutation resulted in 19 months PFS compared to the 5.5 months of those receiving placebo.^3^ Given the impressive success of PARPi in this population of patients, there is great interest in identifying cohorts of BRCA1/2-proficient patients who may be able to benefit from this therapy, as well.^5^

However, even though a large number of patients experience great clinical benefit from PARPi treatment, a significant subpopulation either do not respond or quickly develop resistance to PARPi, despite being predicted to be sensitive based on their BRCA2 status.^3^ In light of this, efforts have been made to understand the roles of individual proteins in mediating resistance to PARPi in the background of BRCA2 deficiency. For example, depletion of RADX or EZH2 has been shown to rescue the cytotoxicity caused by PARP inhibition in BRCA2- deficient cells.^6,7^ Moreover, depletion of PTIP or CHD4 reduces PARPi-induced chromosomal aberrations in BRCA2-deficient cells.^8,9^ However, a comprehensive, genome-wide characterization of potential mediators of PARPi sensitivity and resistance would both advance the fundamental understanding of the processes underlying these effects, as well as potentially promote more effective usage in the clinic.

In order to better understand the mechanisms regulating cellular sensitivity and resistance to PARP inhibitors, we designed complementary genome-wide CRISPR screens in a pair of parental wildtype and BRCA2-knockout (BRCA2^KO^) HeLa cell lines. This approach allowed us to investigate which specific genetic changes lead to PARPi sensitivity in inherently resistant cells (parental) or to resistance in intrinsically sensitive cells (BRCA2^KO^) in an otherwise isogenic background. We validate two proteins previously unconnected to PARPi resistance, namely TIP60 and HUWE1, which cause resistance to PARPi when depleted in BRCA2-deficient cells. Furthermore, we show that the mechanism of resistance caused by TIP60 depletion involves 53BP1-mediated regulation of end resection at double-strand DNA breaks (DSBs).

## Results

### Genome-wide CRISPR screens to identify genetic determinants of the cellular response to the PARPi olaparib

In order to gain a broader understanding of factors governing PARPi response, we performed a series of CRISPR knockout and transcriptional activation (overexpression) screens in an isogenic pair of wildtype and BRCA2-knockout HeLa cell lines.

First, to identify genetic changes which sensitize cells to PARPi treatment, we employed the Brunello human CRISPR knockout pooled library, which targets 19,114 genes with four single-guide RNAs (sgRNAs) per gene.^10^ HeLa cells infected with the Brunello library were divided into PARPi (5 μM olaparib)- or vehicle (DMSO)- treated arms, and after 4 days surviving cells were harvested for sgRNA sequencing and bioinformatic analysis (Figure 1A). By treating parental HeLa cells with a relatively low dose of olaparib and identifying sgRNA sequences which dropped out in the olaparib-treated arm as compared to the DMSO-treated arm, we were able to generate a list of genes which, when knocked out, sensitize wildtype cells to PARPi (Figure 1B, Supplemental Table S1). Importantly, RAD51, an essential component of the homologous recombination pathway, was one of the most significant hits from this screen. In addition, several RAD51 paralogs including RAD51B, XRCC3, and RAD51C were among the top hits; indeed, RAD51C loss via mutation or promoter hypermethylation in tumors has been connected to favorable PARPi response in patients.^11^ Other notable top hits include multiple RNase H2 subunits, consistent with previous findings implicating loss of RNase H2 in sensitizing HR-proficient cells to PARPi.^12^ Collectively, top hits were enriched for biological processes including homologous recombination, DNA replication, translational initiation, and DNA repair (Supplemental Figure S1).

**Figure 1.**
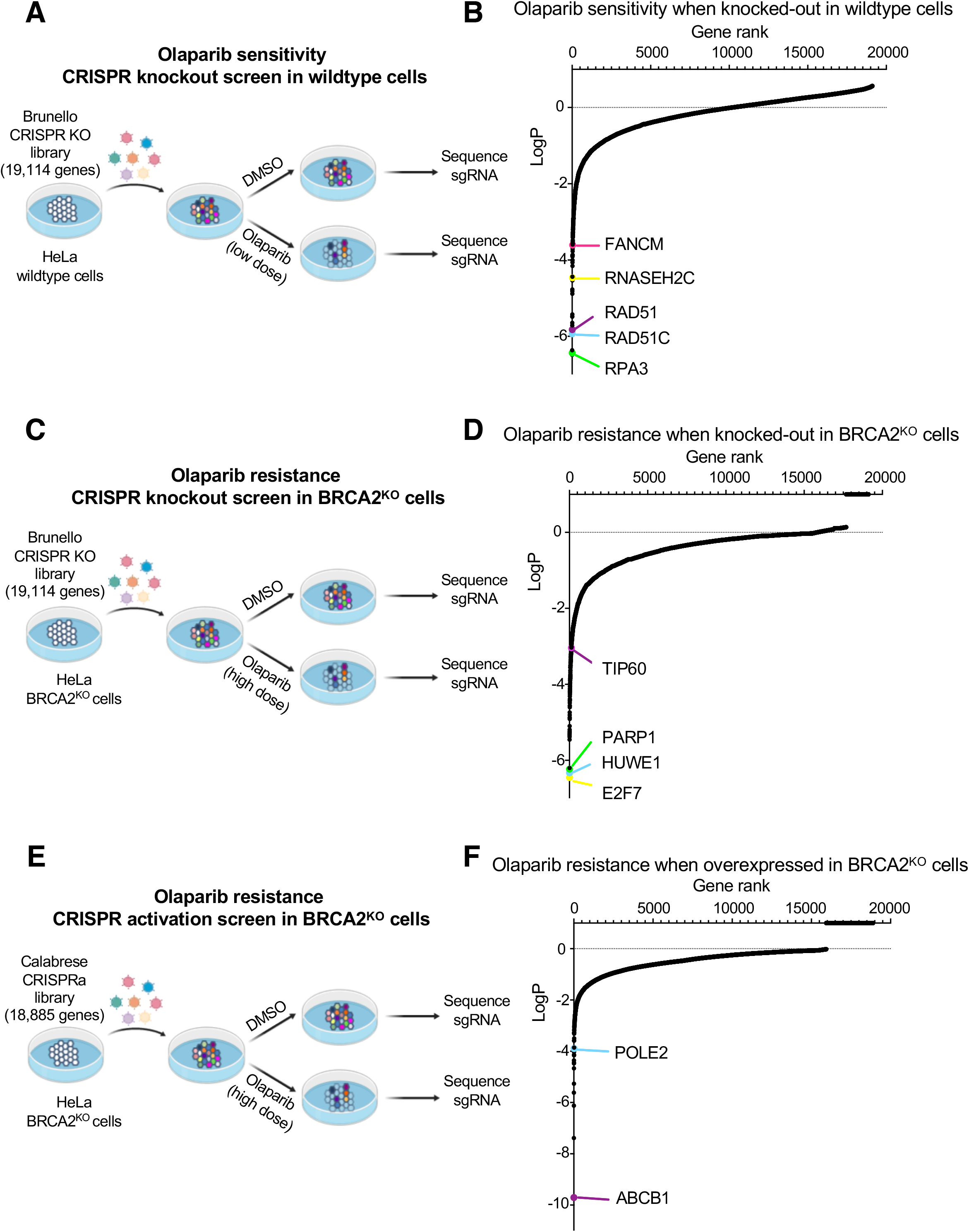
Complementary CRISPR knockout and activation screens identify determinants of PARPi response in parental or BRCA2-knockout HeLa cells. **(A)** Schematic representation of the CRISPR knockout screen for olaparib sensitivity in wildtype cells. HeLa cells were infected with the Brunello CRISPR knockout library. Infected cells were divided into PARP inhibitor (olaparib) -treated or control (DMSO) arms. Genomic DNA was extracted from cells surviving the drug treatment and single-guide RNAs (sgRNAs) were identified using Illumina sequencing. **(B)** Scatterplot showing the results of this screen. Each gene targeted by the library was ranked according to P-values calculated using RSA analysis. The P-values are based on the fold change of the guides targeting each gene between the olaparib- and DMSO-treated conditions. Several biologically interesting hits are highlighted. **(C)** Schematic representation of the CRISPR knockout screen for olaparib resistance in BRCA2^KO^ cells. HeLa BRCA2^KO^ cells were infected with the Brunello CRISPR knockout library. Infected cells were divided into PARP inhibitor (olaparib) -treated or control (DMSO) arms. **(D)** Scatterplot showing the results of this screen, with several biologically interesting hits highlighted. **(E)** Schematic representation of the CRISPR activation screen for olaparib resistance in BRCA2^KO^ cells. HeLa BRCA2^KO^ cells stably expressing the modified dCas9 enzyme were infected with the Calabrese CRISPR activation library. Infected cells were divided into PARP inhibitor (olaparib) -treated or control (DMSO) arms. **(F)** Scatterplot showing the results of this screen, with several biologically interesting hits highlighted.

Next, we sought to identify genes which cause PARPi resistance when depleted in BRCA2^KO^ cells. To identify resistant cells, we treated CRISPR knockout library-infected BRCA2^KO^ cells with a dose olaparib (4 μM) which killed more than 90% of cells over 4 days (Figure 1C). Then, we searched for sgRNA sequences which were enriched in the cells surviving olaparib treatment as compared to DMSO (Figure 1D, Supplemental Table S1). It has been well-established in previous studies that PARP1 itself is required for the cytotoxicity of PARPi.^13–15^ In line with this, we found that PARP1 was ranked among the most significant of hits in our screen, as well (Figure 1D). Another top hit from our screen was the transcription factor E2F7 (Figure 1D). We previously showed that loss of E2F7 causes PARPi resistance in BRCA2-deficient cells through transcriptionally upregulating RAD51 and subsequently restoring homologous recombination.^16^ Unlike as seen in the sensitivity screen, the pathway analysis of the resistance screen implicated more unexpected pathways such as vesicle-mediated transport and tRNA processing in PARPi resistance (Supplemental Figure S2). While this finding could reflect a role for these pathways in PARPi resistance, an alternative explanation may be that proteins in these pathways have heretofore underappreciated roles in DNA repair.

Finally, we performed a CRISPR activation screen to identify genes which cause PARPi resistance when overexpressed in BRCA2^KO^ cells. We utilized the Calabrese human CRISPR activation library, which transcriptionally activates 18,885 genes individually by using guide RNA sequences to recruit an enzymatically dead Cas9 (dCas9) and transcriptional activators to the region near the transcriptional start site of the gene.^17^ Screening conditions were maintained as performed in the resistance CRISPR knockout screen. Briefly, HeLa BRCA2^KO^ cells expressing dCas9 were infected with the activation library and divided into high dose PARPi (4 μM olaparib)- or vehicle (DMSO)- treated arms (Figure 1E, Supplemental Figure S3A). Roughly 90% of cells were killed after 4 days of treatment; surviving cells were harvested and analyzed to identify genes which cause resistance to PARPi when transcriptionally activated (Figure 1F, Supplemental Figure S3B, Supplemental Table S1). There are few studies reporting proteins which cause PARPi resistance when overexpressed; however, interestingly, our top hit from the screen, ABCB1, has previously been identified as a mechanism of PARPi resistance.^18^ This gene encodes the protein MDR-1 (multidrug resistance protein 1), a drug efflux pump whose overexpression was associated with acquired resistance to olaparib in ovarian cancer cell lines.^18^ To validate these results in our system, we generated HeLa BRCA2^KO^ cells with transcriptional activation of the ABCB1 gene (Figure 2A). We found that the BRCA2-knockout, ABCB1-overexpressing cell line was as resistant to PARPi as the BRCA2-proficient parental HeLa line (Figure 2B). Previously, PARPi treatment has been shown to induce apoptosis in BRCA2-deficient cells.^19^ In line with this, we found that olaparib treatment led to a more than 6-fold increase in cells positive for annexin V, a marker of apoptosis, in BRCA2-knockout cells; however, overexpression of ABCB1 restored olaparib-induced apoptosis to control levels (Figure 2C). Overall, these findings validate the results of our CRISPR activation screen.

**Figure 2.**
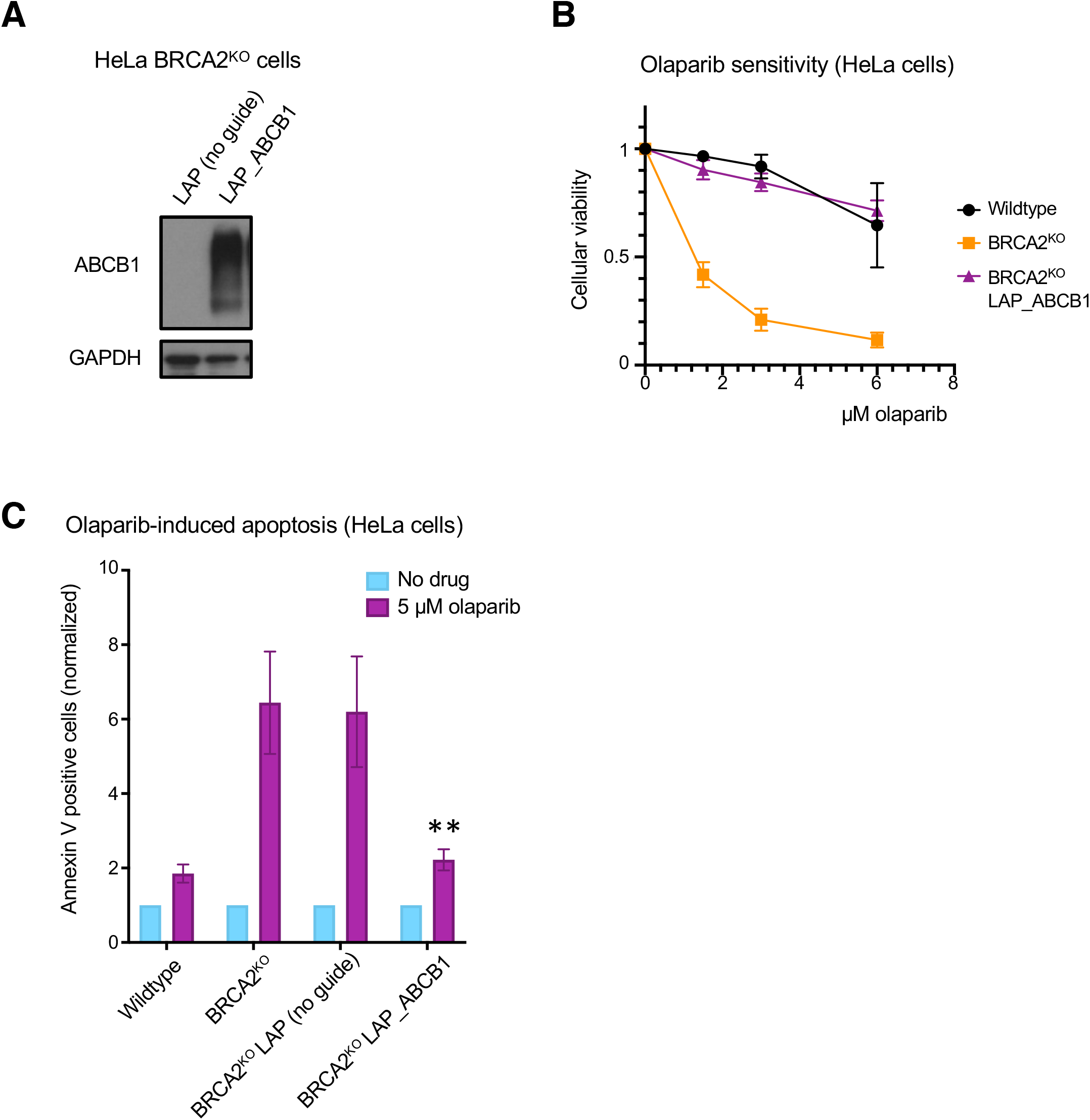
Overexpression of ABCB1, the top hit from the CRISPR activation screen, causes PARPi resistance in BRCA2-deficient cells. **(A)** Western blot showing overexpression of ABCB1 in the cell line containing all three components of the CRISPR lentiviral activation particle (LAP) system (dCas9, activator helper complex, and sgRNA targeting the ABCB1 gene) but not in the control cell line lacking the sgRNA. **(B)** Cellular viability assay showing that ABCB1 transcriptional activation rescues PARPi sensitivity in HeLa BRCA2-knockout cells. The averages of 4 experiments are shown, with standard deviations as error bars. **(C)** Olaparib-induced apoptosis in BRCA2-knockout cells is suppressed by ABCB1 overexpression. The averages of 4 experiments are shown, with standard deviations as error bars. Asterisks indicate statistical significance (compared to “no guide” sample).

### Depletion of TIP60 or HUWE1 rescues PARPi sensitivity in BRCA2-deficient cells

From the PARPi resistance screen in BRCA2^KO^ cells, two DNA repair genes previously unconnected to PARPi resistance, namely the histone acetyltransferase TIP60 (KAT5) and E3 ubiquitin ligase HUWE1, drew our attention. We first sought to validate these hits. In order to test the effect of TIP60 or HUWE1 loss on PARPi sensitivity in BRCA2-depleted cells, we knocked down these genes and assessed cellular viability after olaparib treatment. Strikingly, TIP60 or HUWE1 knockdown in HeLa cells depleted of BRCA2 using siRNA led to PARPi resistance similar to that seen in HR-proficient controls (Figure 3A). This robust rescue of PARPi sensitivity was also observed in our HeLa BRCA2^KO^ cells upon depletion of TIP60 (Supplemental Figures S4A and S4B) or HUWE1 (Supplemental Figures S4C and S4D) using either of two siRNA targeting sequences. We next asked whether these phenotypes were cell line-specific by extending our studies into additional cell lines. While TIP60 or HUWE1 depletion alone did not affect sensitivity of wildtype controls to PARPi, co-depletion of either protein in BRCA2-depleted U2OS (Figure 3B) or BRCA2-knockout DLD1 (Figure 3C) cells significantly reduced PARPi sensitivity.

**Figure 3.**
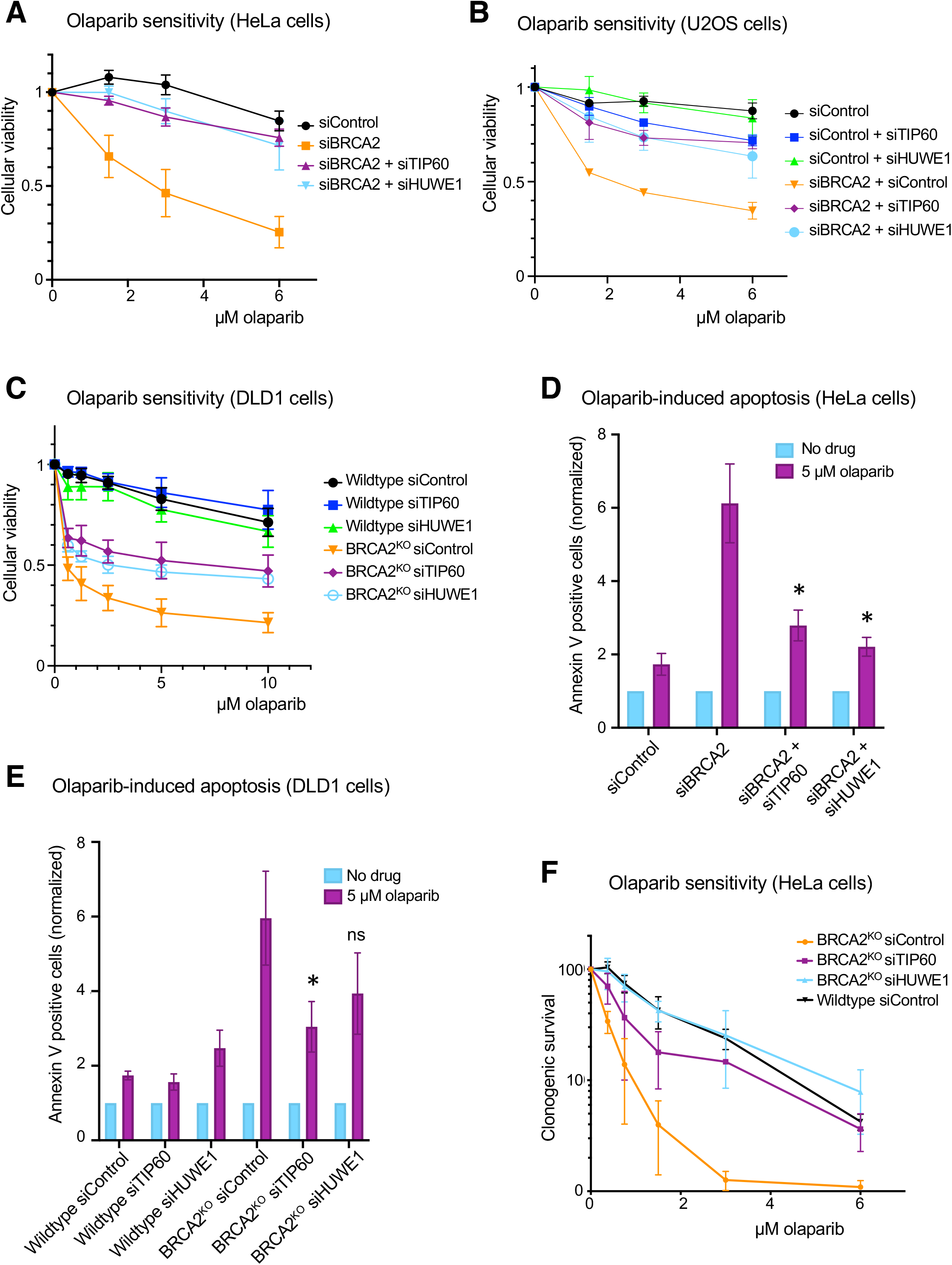
TIP60 or HUWE1 knockdown in BRCA2-depleted cells results in PARPi resistance. **(A-C)** Knockdown of TIP60 or HUWE1 rescues the olaparib sensitivity of BRCA2-knockdown HeLa **(A)**, BRCA2-knockdown U2OS **(B)**, and BRCA2-knockout DLD1 **(C)** cells in cellular viability assays. **(D, E)** Olaparib-induced apoptosis is rescued by TIP60 or HUWE1 depletion in BRCA2-knockdown HeLa (**D**) and BRCA2-knockout DLD1 (**E**) cells. Asterisks indicate statistical significance, compared to siBRCA2 (**D**) or BRCA2^KO^ siControl (**E**) samples. **(F)** TIP60 or HUWE1 depletion rescues sensitivity of BRCA2-knockout HeLa cells to PARPi in a clonogenic survival assay. All figures reflect the average of three experiments performed after 72 hours of treatment, with standard deviations shown as error bars.

We next investigated the effect of TIP60 or HUWE1 depletion on olaparib-induced apoptosis using Annexin V flow cytometry. Consistent with the results of the cellular viability assays, depletion of TIP60 or HUWE1 abrogated the increase in Annexin V–positive cells caused by PARPi treatment in BRCA2-knockdown HeLa cells as well as BRCA2-knockout DLD1 cells (Figures 3D and 3E). We observed similar phenotypes using a second siRNA oligonucleotide for each hit in HeLa BRCA2^KO^ cells (Supplemental Figures S4E and S4F).

In light of this increased cellular survival after PARPi treatment in BRCA2-deficient cells depleted of the two hits, we wondered if these surviving cells would demonstrate improved viability in the long-term. To test this, we performed clonogenic survival assays in HeLa BRCA2^KO^ cells. Cells were pre-treated with siRNAs targeting TIP60, HUWE1, or a control siRNA, treated with olaparib for three days, and then allowed to form colonies for two weeks in drug-free media. We found that while BRCA2^KO^ cells showed reduced colony-forming ability after olaparib treatment, TIP60- and HUWE1- depleted BRCA2^KO^ cells were able to form colonies after olaparib treatment in a manner similar to wildtype controls (Figure 3F). Taken together, these findings confirm the results of the CRISPR knockout screen and demonstrate that TIP60 or HUWE1 depletion leads to PARPi resistance in BRCA2-deficient cells.

### PARPi resistance caused by TIP60 depletion is dependent on the 53BP1/REV7 pathway

After observing that TIP60 depletion robustly rescued PARPi sensitivity, we sought to investigate the mechanisms through which this may occur. To this end, we tested the effect of TIP60 depletion on several previously proposed mechanisms of PARPi-induced cytotoxicity. Trapping of PARP1 on the chromatin has been suggested as an underlying cause of the toxicity of PARPi.^13^ Given that TIP60 has been previously connected to the flux of PARP1 on chromatin,^20^ we tested if TIP60 depletion may abrogate this PARP-trapping effect of the inhibitor. However, we observed no difference in trapped PARP1 after TIP60 depletion using a chromatin fractionation assay (Supplemental Figure S5A). Furthermore, an aberrant increase in replication fork speed was recently proposed as a mechanism of action of PARPi.^21^ While we did observe an increase in replication fork speed upon PARPi treatment, this phenotype was not affected by TIP60 depletion (Supplemental Figure S5B). Finally, we have previously shown that E2F7 depletion led to PARPi resistance in BRCA2-deficient cells by rescuing the defect in homologous recombination.^16^ In contrast, TIP60 depletion did not improve the HR efficiency of BRCA2-depleted cells (Supplemental Figure S5C). Interestingly, however, despite no change in HR repair, we observed a reduction in olaparib-induced double-strand breaks in HeLa BRCA2^KO^ cells after TIP60 depletion using a neutral comet assay (Figure 4A), suggesting that repair of DSBs is improved in these cells.

**Figure 4.**
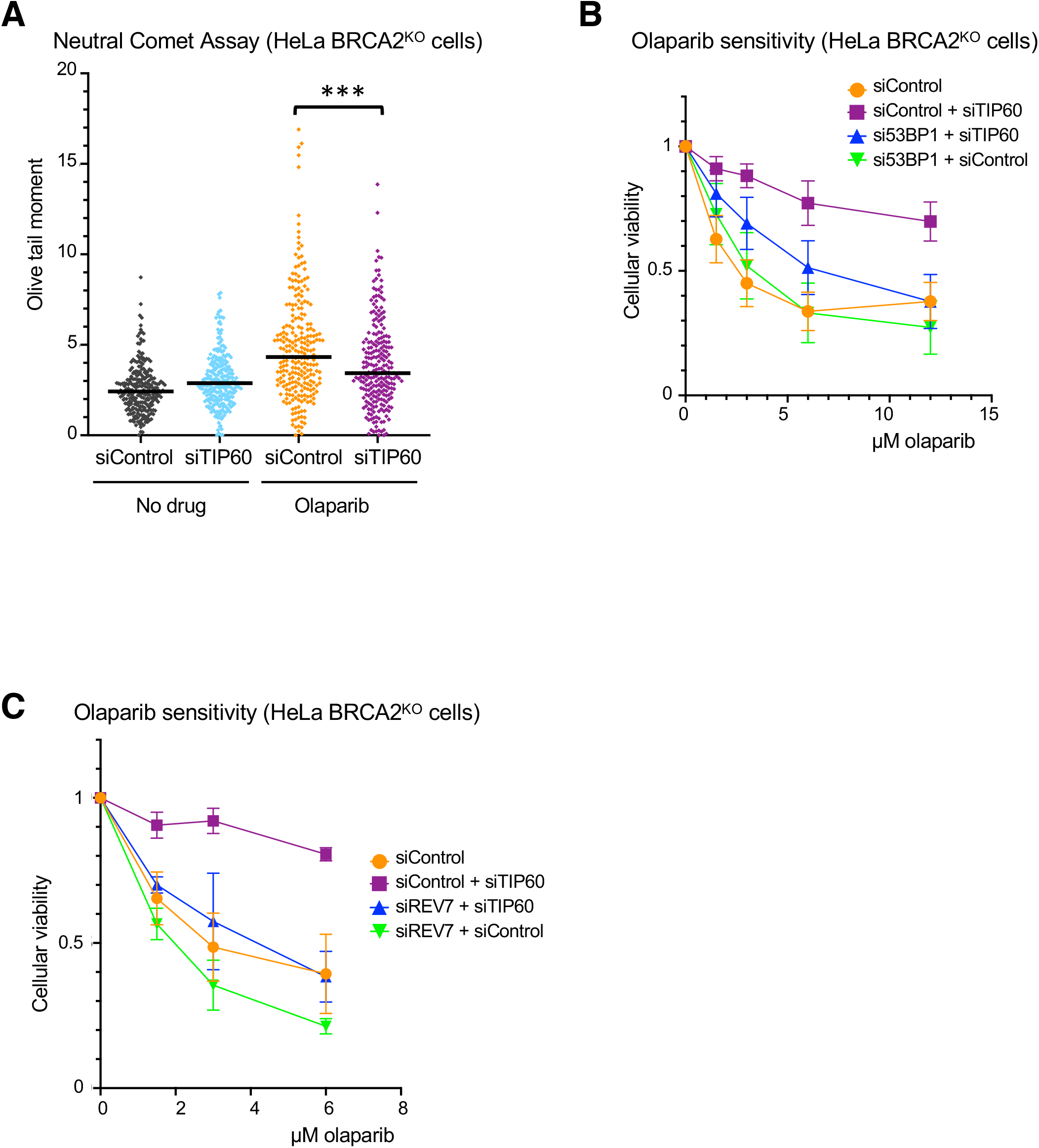
TIP60 depletion reduces olaparib-induced double-strand breaks and relies on the 53BP1/REV7 pathway to rescue olaparib-induced cytotoxicity. **(A)** Neutral Comet assay showing that olaparib treatment (10 μM for 24 hours) induces double-strand breaks in HeLa BRCA2-knockout cells, and that this effect is abrogated by TIP60 depletion. Approximately 120 comets from each of two experiments were pooled for each sample. Medians are shown as horizontal bars. Asterisks indicate statistical significance. **(B)** PARPi resistance caused by TIP60 depletion is dependent on 53BP1. The averages of 9 experiments are shown, with standard deviations as error bars. **(C)** REV7 is required for the PARPi resistance after TIP60 knockdown. The averages of 3 experiments are shown, with standard deviations as error bars.

Previous studies have indicated that TIP60 activity antagonizes the recruitment of 53BP1 to damaged DNA.^22–24^ 53BP1 is a key mediator of double-strand break repair pathway choice, which suppresses DNA end resection at DSBs and promotes non-homologous end joining (NHEJ).^25,26^ We reasoned that increased 53BP1 binding due to TIP60 depletion may be involved in the improved cellular viability and reduced double-strand breaks observed after olaparib treatment in BRCA2-deficient cells. To test if 53BP1 is required for PARPi resistance in BRCA2-deficient cells depleted of TIP60, we performed additional cellular viability assays. TIP60 was depleted alone or in combination with 53BP1 in HeLa BRCA2^KO^ cells and sensitivity to PARPi was assessed. We found that while TIP60 depletion alone causes resistance, co-depletion of 53BP1 with TIP60 severely diminishes this rescue (Figure 4B). Importantly, the magnitude of TIP60 knockdown was unaffected by 53BP1 co-depletion (Supplemental Figure S6A). Recently, the Shieldin complex has been identified as a downstream effector of 53BP1 at the double-strand break.^27,28^ Therefore, we next tested if REV7, a component of the Shieldin complex, was also required for the rescue of PARPi sensitivity caused by TIP60 loss. Indeed, co-depletion of REV7 abolished the PARPi resistance produced by the depletion of TIP60 (Figure 4C and Supplemental Figure S6B). Taken together, these results demonstrate that the 53BP1/REV7 pathway is required for PARPi resistance caused by TIP60 depletion in BRCA2-deficient cells.

### TIP60 depletion increases 53BP1 binding at double-strand breaks and reduces end resection

We next sought to directly assess the impact of TIP60 depletion on the binding of 53BP1 near the double-strand break site in BRCA2-deficient cells. We used a chromatin immunoprecipitation (ChIP) assay with a 53BP1 antibody to evaluate binding near the site of an inducible double-strand break within an integrated reporter in the previously described U2OS-DSB reporter cell line.^24^ qPCR analysis of the immunoprecipitated DNA revealed increased 53BP1 binding near the double-strand break site after TIP60 knockdown in BRCA2-depleted cells (Figure 5A).

**Figure 5.**
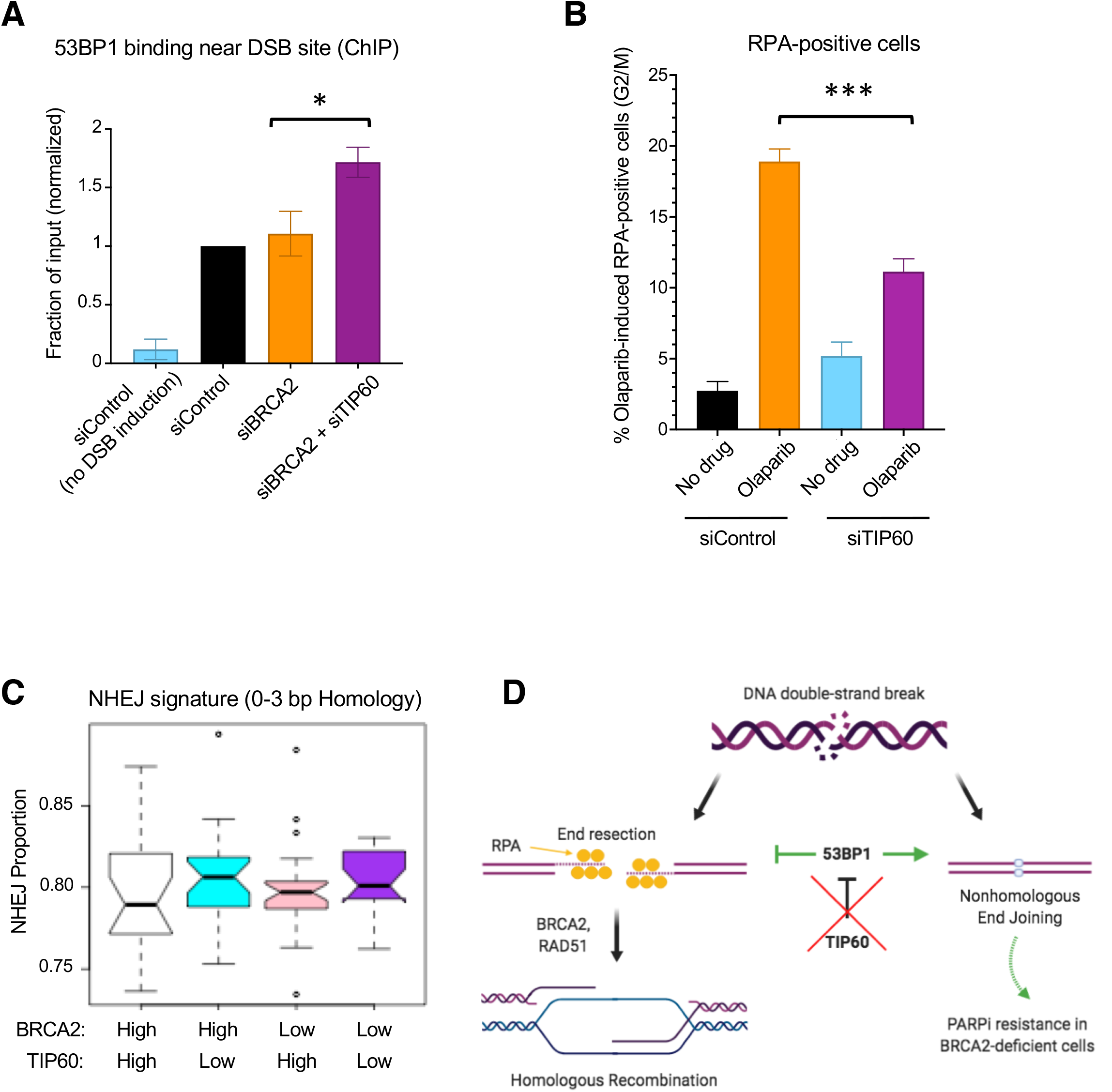
Functional consequences of TIP60 depletion. **(A)** Depletion of TIP60 leads to increased recruitment of 53BP1 near the site of the double-strand break in BRCA2-depleted U2OS-DSB reporter cells. DSBs were generated at a specific genomic locus through induction of expression of LacI-FokI nuclease. Then, chromatin immunoprecipitation was performed with an antibody against 53BP1 and qPCR was performed using primers for the DSB site. The averages of 3 experiments are shown, with standard error of the mean as error bars. Asterisks indicate statistical significance from a paired TTEST. **(B)** TIP60 loss leads to a reduction in olaparib-induced end resection as measured by quantification of RPA-positive cells. Cells were treated with 10 μM olaparib for 12 hours before being harvested for flow cytometry experiments using an RPA antibody. Representative gating of these experiments is shown in Supplemental Figure S6C. The averages of 3 experiments are shown, with standard deviations as error bars. Asterisks indicate statistical significance. **(C)** Bioinformatic analysis of the OV-AU dataset reveals that TIP60 low tumors show increased NHEJ signature at structural variations breakpoints, as compared to TIP60 high tumors, in both BRCA2 high and BRCA2 low backgrounds. **(D)** Proposed model for resistance caused by TIP60 depletion. In BRCA2-proficient cells, homologous recombination is the most accurate mechanism of double-strand break repair. However, this pathway is not functional in BRCA2-deficient cells. TIP60 depletion increases 53BP1 binding at the double-strand break in BRCA2-deficient cells, preventing the break from proceeding down the “dead-end” process of HR and instead directs it towards repair by NHEJ.

As the 53BP1/Shieldin pathway functions to prevent excessive end resection at DNA ends, we sought to investigate the effect of TIP60 depletion on end resection.^28^ To test this, we quantified the percentage of cells that were positive for RPA, a protein which binds to single-stranded DNA produced by end resection, using flow cytometry as previously described.^29,30^ We observed an accumulation of S-phase cells in control cells treated with olaparib, which was not observed in siTIP60-treated cells (Supplemental Figure S6C). To prevent these differences from confounding our analysis, we only assessed RPA-positivity in G2 and M -phase cells. In this population of cells, olaparib treatment caused an increase in the percentage of RPA-positive cells, which was reduced upon TIP60 depletion (Figure 5B). Overall, these findings indicate that TIP60 depletion increases 53BP1 binding to olaparib-induced DSBs in BRCA2-deficient cells, suppressing end resection.

Altogether, we observed that TIP60 depletion causes PARPi resistance in a manner dependent on the 53BP1/REV7 pathway, and subsequently causes a reduction in olaparib-induced end resection. Notably, inhibiting end resection directly via depletion of CTIP was not sufficient to cause PARPi resistance in BRCA2-deficient cells (Supplemental Figure S6D and S6E). Previously, exhaustion of 53BP1 by an abundance of DSBs (as one might expect in BRCA2-deficient cells treated with PARPi) was shown to lead to an increase in single strand annealing (SSA), a repair pathway downstream of end resection that results in large deletions and genomic instability.^31^ Therefore, we reasoned that the observed increase in 53BP1 binding may rescue PARPi sensitivity by preventing SSA. However, inhibiting SSA via depletion of LIG1 also failed to cause PARPi resistance in this context (Supplemental Figure S6D and S6E). One potential explanation for these findings is that increased 53BP1 recruitment not only reduces end resection, but also promotes DSB repair through NHEJ.^26^ To address this, we investigated NHEJ DNA repair signature at somatic structural variation breakpoints in publicly available genomic datasets. Bioinformatic analysis of the Australian Ovarian Cancer Study cohort (OV-AU) dataset suggests a higher incidence of NHEJ in ovarian tumors with low TIP60 expression, in both BRCA2-high and BRCA2-low backgrounds (Figure 5C). Overall, this leads to a model in which TIP60 depletion increases 53BP1 binding near the double-strand break, leading to a reduction in end resection and subsequent PARPi resistance, potentially through an increase in NHEJ (Figure 5D).

### TIP60 levels impact cisplatin resistance

Several factors which confer resistance to PARPi have also been shown to rescue sensitivity to the DNA damaging agent and widely used chemotherapeutic, cisplatin.^32^ Thus, we reasoned that TIP60 depletion may also confer resistance to cisplatin in BRCA2-deficient cells. Using a clonogenic survival assay, we found that depletion of TIP60 in BRCA2-knockout HeLa cells rescues cisplatin sensitivity back to that of wildtype cells (Figure 6A). Unlike PARPi, which are relatively new agents in the clinic, cisplatin has remained a mainstay of ovarian cancer treatment for decades.^33^ Thus, we next sought to investigate if TIP60 expression impacts survival of BRCA2-mutant ovarian cancer patients by mining survival data and matched genotype and expression data from publicly available datasets. We queried the OV-TCGA database and asked if ovarian cancer patients with low or high TIP60 expression showed any differences in survival, in BRCA2-mutant or BRCA2-wildtype backgrounds. We found that low TIP60 expression in tumors trended towards poorer survival in patients harboring BRCA2 mutations, but not in the BRCA2-wildtype cohort (Figure 6B). As it is highly likely that most patients in this dataset have been treated with cisplatin, these findings are consistent with increased therapy resistance in the TIP60-low BRCA2 mutant group. Taken together, these data indicate that TIP60 depletion confers resistance to cisplatin in BRCA2-deficient cells and potentially also in tumors.

**Figure 6.**
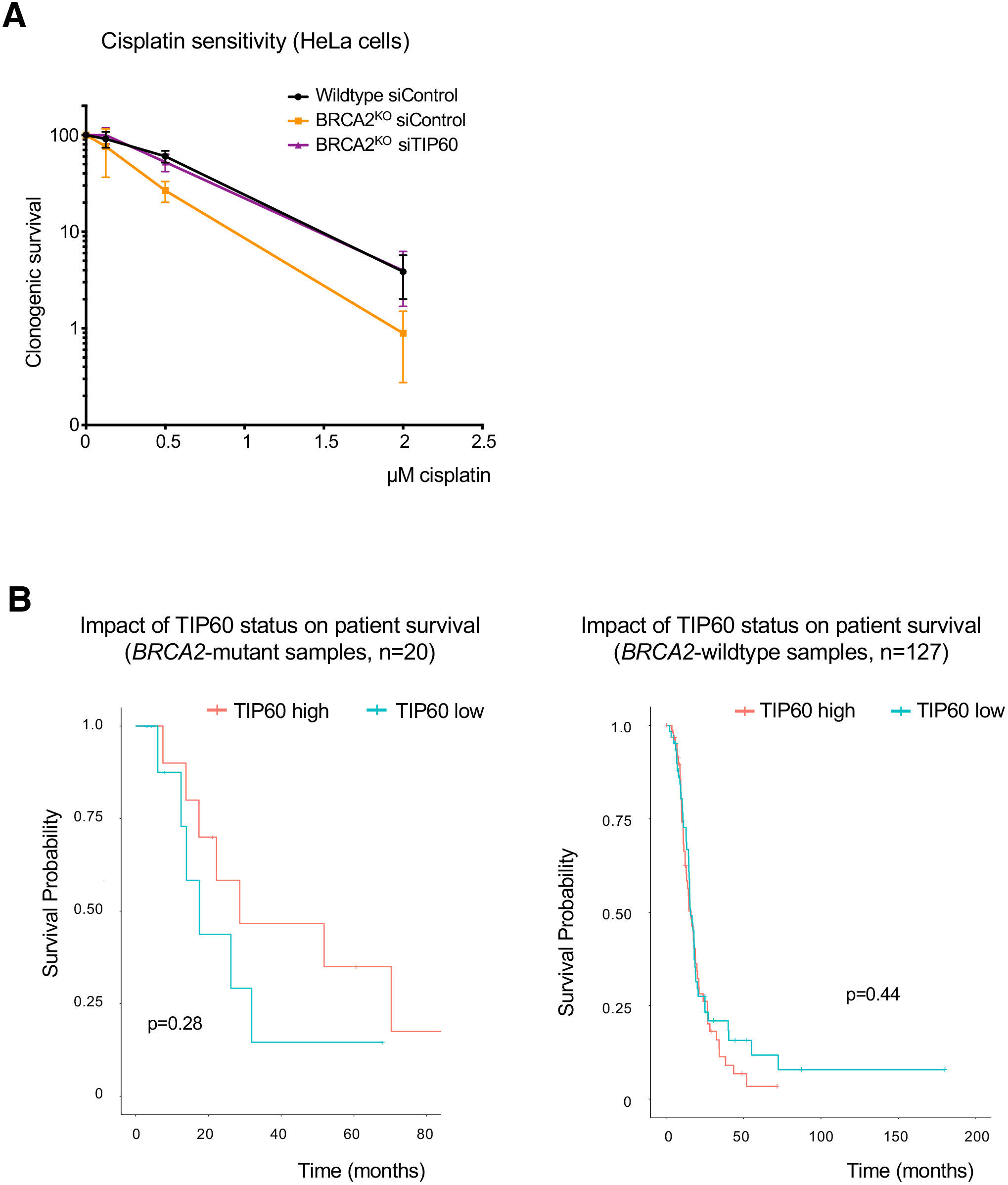
Loss of TIP60 expression causes resistance to cisplatin in BRCA2-deficient cells. **(A)** In clonogenic survival assays, depletion of TIP60 in BRCA2-knockout HeLa cells led to increased colony formation after cisplatin treatment (24 hours). The averages of 3 experiments are shown, with standard deviations as error bars. **(B)** Bioinformatic analysis of the OV-TCGA dataset shows that patients with tumors that have low TIP60 expression trend towards poorer survival in BRCA2 mutant (left) but not BRCA2 wildtype (right) tumors. The difference in BRCA2 mutant samples is not significant (p=0.28) likely because of the small number (n=20) of BRCA2-mutant samples in this dataset.

## Discussion

Identifying potential predictors of response to PARPi serves to both identify additional patients who may benefit from this therapy, and to avoid ineffective treatment for those who will not. Additionally, beyond simply predicting treatment response, achieving a better understanding of mechanisms of sensitivity and resistance to PARPi brings us closer to potential therapeutic interventions that may be able to sensitize or re-sensitize patients to treatment. Therefore, there has been great interest in investigating PARPi sensitivity and resistance through genome-wide CRISPR knockout screens in cells with various genetic backgrounds.^12,32,34^

Here, we have completed a series of genome-wide screens based on CRISPR technology as an unbiased approach towards better understanding determinants of PARPi response. First, we present data from a CRISPR-knockout screen in wildtype HeLa cells, which identified genes which, when depleted, lead to PARPi sensitivity. Additionally, we investigated PARPi resistance in BRCA2-knockout HeLa cells by identifying genes which cause resistance when depleted (CRISPR knockout library) or overexpressed (CRISPR activation library). By utilizing two different types of libraries and otherwise isogenic cell lines differing only in BRCA2 levels, we are able to not only examine the results of each screen individually, but also to consider the results in relationship to one another. For example, loss of POLE3 or POLE4, subunits of DNA polymerase Epsilon, sensitized parental HeLa cells to PARPi treatment, while transcriptional activation of polymerase Epsilon subunit POLE2 was associated with PARPi resistance in BRCA2-knockout HeLa cells. This is especially interesting given that depletion of polymerase Epsilon subunits was recently shown to lead to sensitivity to ATR inhibitors.^35^

Our results identified several hits which were consistent with previous findings from other groups, as well as novel hits which have not previously been connected to PARPi response. One such novel hit is TIP60, which we validate using several cellular survival and apoptosis assays, in multiple cell lines. Furthermore, we show that this rescue of PARPi-induced cytotoxicity is dependent on both 53BP1 and REV7. Functionally, TIP60 depletion increases the binding of 53BP1 near the double-strand break, reduces olaparib-induced end resection, and ameliorates the olaparib-induced increase in double-strand breaks. This is a novel mechanism of PARPi resistance in BRCA2-deficient cells, and highlights critical differences between BRCA2- and BRCA1-deficiency. In fact, whereas we show that in BRCA2-deficient cells an increase in 53BP1 binding to damaged DNA plays a role in PARPi resistance, others have shown 53BP1 loss to be a mechanism of PARPi resistance in BRCA1-deficent cells.^25,36^ This divergent impact of 53BP1 on PARPi resistance in BRCA2- and BRCA1- deficient cells likely reflects the different roles of BRCA1/2 in the double-strand break repair process of homologous recombination, and is consistent with previous reports that 53BP1 depletion rescued defects in proliferation and checkpoint responses in BRCA1-deficient, but not BRCA2-deficient cells.^37^ BRCA1 functions early in homologous recombination, promoting end resection at the double-strand break; this is in direct opposition to the anti-resection role of 53BP1.^26^ Therefore, 53BP1 depletion would be expected to alleviate the defects caused by BRCA1 deficiency and improve the ability of the cell to manage olaparib-induced damage, resulting in resistance. On the other hand, BRCA2 functions much later in the process of homologous recombination. Hence, 53BP1 depletion would not correct this defect; instead, increased 53BP1 binding would reduce end resection and prevent the break from beginning down the homologous recombination pathway, which is a “dead-end” in BRCA2-deficient cells. Altogether, our work identifies a novel mechanism of PARPi resistance in BRCA2-deficient cells and emphasizes the fact that despite both causing defects in homologous recombination, BRCA1- and BRCA2- deficiency represent distinct defects that should not always be grouped together in the laboratory or the clinic.

We show here that 53BP1 is required for the rescue of PARPi sensitivity caused by TIP60 depletion, and also that TIP60-deficient patient tumors show an increase in the non-homologous end joining pathway of double-strand break repair, consistent with the expected effects of an increase in 53BP1 binding at the double-strand break.^25,26^ However, 53BP1 has been implicated in additional PARPi-relevant contexts, such as at the replication fork.^38–40^ The finding that REV7 is also required for the resistance caused by TIP60 depletion supports a mechanism involving activity at the double-strand break;^41^ however, other potential roles for REV7 at the replication fork have not yet been thoroughly characterized.^42^ Therefore, other function(s) of 53BP1/REV7 may potentially be involved in the rescue of PARPi-induced cytotoxicity downstream of TIP60 depletion in BRCA2-deficient cells.

## Materials and methods

### Cell culture

Human HeLa, U2OS, and RPE cells were grown in Dulbecco’s modified Eagle’s medium (DMEM). DLD1 cells were grown in Roswell Park Memorial Institute (RPMI) 1640 medium. DMEM and RPMI were supplemented with 10% fetal bovine serum. HeLa BRCA2^KO^ cells were generated in our laboratory and previously described.^16^ DLD1 BRCA2^KO^ cells (Horizon HD105-007) were obtained from Dr. Robert Brosh (National Institute on Aging, NIH). U2OS-DSB reporter cells were obtained from Dr. Roger Greenberg (University of Pennsylvania).^24^ U2OS DR-GFP cells were obtained from Dr. Jeremy Stark (City of Hope National Medical Center, Duarte, CA).^43^

Gene knockdowns were performed using Lipofectamine RNAiMAX reagent for transfection of Stealth siRNA (Life Tech, unless otherwise noted). Oligonucleotide sequences used were:

BRCA2: GAGAGGCCTGTAAAGACCTTGAATT
TIP60 #1: GATGGACGTAAGAACAAGAGTTATT
TIP60 #2: CACCCATTCATCCAGACGTTTGTTG
HUWE1 #1: TTTAAGGGTGGGCTGATGTCTCATG
HUWE1 #2: CACACCAGCAATGGCTGCCAGAATT
53BP1: TCCCAGAGTTGATGTTTCTTGTGAA
REV7: GTGGAAGAGCGCGCTCATAAA (Qiagen)
CTIP: GGGTCTGAAGTGAACAAGATCATTA
LIG1: CCAAGAACAACTATCATCCCGTGGA

For all genes, siRNA #1 was used for knockdown unless otherwise indicated. Knockdown was confirmed by western blot for all proteins targeted, except REV7, which was assessed using real-time quantitative PCR (RT-qPCR) as previously described.^16^ Primers used were:

REV7 for: TGCTGTCCATCAGCTCAGAC;
REV7 rev: TCTTCTCCATGTTGCGAGTG;
GAPDH for: AAATCAAGTGGGGCGATGCTG;
GAPDH rev: GCAGAGATGATGACCCTTTTG.

HeLa BRCA2-knockout cells overexpressing ABCB1 via transcriptional activation were created by consecutive rounds of transduction and selection. Cells were first infected with dCas9 (Addgene, 61425-LV) and colonies were selected with blasticidin, 3 μg/ml. dCas9-expressing cells were then infected with the MS2-P65-HSF1 activator helper complex (Addgene, 61426-LVC) and treated with 0.5 mg/ml hygromycin. Finally, hygromycin-resistant cells were infected with lentivirus containing guide RNA targeting ABCB1 (Sigma Custom CRISPR in lentiviral backbone LV06), guide sequence (5’-3’): GGGAGCAGTCATCTGTGGTG. Infected cells were selected using puromycin (0.6 μg/ml).

### Genome-wide CRISPR screens

For the CRISPR knockout screens, wildtype or BRCA2-knockout HeLa cells were transduced with the Brunello Human CRISPR knockout pooled library (Addgene, 73179).^10^ To achieve a representation of 250 cells per sgRNA, 50 million cells were transduced at a low multiplicity of infection (MOI) (0.4). Selection with puromycin (0.6 μg/ml) was initiated at 48 hours post-transduction and maintained for 4-6 days. For the resistance screen in BRCA2-deficient cells, puromycin-resistant cells were divided into olaparib- or vehicle- (DMSO) treated arms at 250 fold library coverage per arm. Olaparib was used at 4μM for 4 days, and yielded 7% survival relative to the DMSO condition. After treatment, cells were pelleted and flash-frozen for DNA extraction. For the sensitivity screen in wildtype cells, puromycin-resistant cells were seeded at a representation of 150 cells per sgRNA and treated with DMSO or olaparib (5μM) for 4 days. After the 4 days of treatment, survival of the olaparib-treated population was 43% of the vehicle-treated cells; cells were pelleted and flash-frozen for DNA extraction.

#### CRISPR Activation Screen

For the CRISPR activation screen, HeLa BRCA2-knockout cells were infected with dCas9 (Addgene, 61425-LV) and selected with blasticidin (3 μg/ml). dCas9-expressing cells were then transduced with the Calabrese Human CRISPR Activation Pooled Library (Set A, Addgene, 92379-LV) using enough cells to obtain a library coverage of 500 cells per sgRNA at an MOI of 0.4.^17^ Selection with puromycin (0.6 μg/ml) was initiated at 48 hours post-transduction and maintained for 5 days. After puromycin selection, cells were divided into olaparib- (4 μM) or vehicle- (DMSO) treated arms at 500-fold library coverage per arm. After 4 days of treatment, survival of the olaparib-treated cells was 12% relative to the DMSO-treated condition; cells were flash-frozen as pellets for DNA extraction.

### Sequencing and analysis of CRISPR screens

Genomic DNA (gDNA) was extracted per manufacturer’s instructions using the DNeasy Blood & Tissue Kit (Qiagen, 69504). A maximum of 5 million cells were used per column. Isolated gDNA was quantified using Nanodrop. The genomic DNA equivalent of 125-fold library coverage (knockout screens) or 250-fold library coverage (activation screen) was used as template for PCR amplification of the sgRNA sequences. Each PCR reaction included no more than 10 μg of gDNA, in addition to the following components: 3 μl of Radiant HiFi Ultra Polymerase (Stellar Scientific, RAD-HF1100), 20 μl of 5X HiFi Ultra Reaction Buffer, 4 μl of 10 μM P5 primer, 4 μl of 10 μM uniquely-barcoded P7 primer, and water to bring the total reaction volume to 100 μl. Primers were synthesized by Eurofins Genomics using the sequences listed in the user guide provided for the CRISPR libraries. PCR cycling conditions were as follows: an initial 2 min at 98°C; followed by 10 s at 98°C, 15 s at 60°C, 45 s at 72°C, for 30 cycles; and lastly 5 min at 72°C.^44^ The E.Z.N.A. Cycle Pure Kit (Omega, D6493-02) was used per manufacturer’s instructions to purify PCR products. PCR products were further purified with Agencourt AMPure XP SPRI beads according to manufacturer’s instructions (Beckman Coulter, A63880). The final product was assessed for its size distribution and concentration using BioAnalyzer High Sensitivity DNA Kit (Agilent Technologies) and qPCR (Kapa Biosystems). Pooled libraries were diluted to 2 nM in EB buffer (Qiagen) and then denatured using the Illumina protocol. The denatured libraries were diluted to 10 pM by pre-chilled hybridization buffer and loaded onto a TruSeq v2 Rapid flow cell on an Illumina HiSeq 2500 and run for 50 cycles using a single-read recipe according to the manufacturer’s instructions (Illumina).

For bioinformatic analysis, de-multiplexed and adapter-trimmed sequencing reads were generated using Illumina bcl2fastq (released version 2.18.0.12) allowing no mismatches in the index read. Analysis was then performed following a previously described protocol.^45^ Briefly, sgRNA representation was analyzed for each condition using the provided custom python script (count_spacers.py) (step 38^45^). The sgRNA fold change due to olaparib treatment was then determined for each cell line as follows (Step 65^45^): A pseudocount of 1 was added to each sgRNA read count, which was then normalized by the total reads for that condition; the normalized sgRNA read counts of the olaparib-treated condition were then divided by those of the DMSO-treated condition, and the base 2 logarithm of the resulting ratios were calculated. These values were then used as inputs for analysis using the redundant siRNA activity (RSA) method.^46^ For the analysis, the Bonferroni correction option was used. Bounds were adjusted such that for the sensitivity screen, guides which were 2-fold less present in the olaparib treated condition were counted as guaranteed hits. For the resistance screens the default bounds were used, in which guides which were 2-fold more enriched in cells surviving olaparib treatment were counted as guaranteed hits.

### Protein techniques

Total cellular protein extracts and western blots were performed as previously described.^47^ For the PARP trapping assay, cells were co-treated with MMS (0.01%) and olaparib (1μM) for 3 hours to induce trapping of PARP1 as previously described.^13^ Cellular fractionation was performed with the Subcellular Protein Fractionation Kit for Cultured Cells (Thermo Scientific, 78840) per manufacturer’s instructions; protein was quantified using the Qubit Protein Assay Kit (Invitrogen). Antibodies used were: ABCB1 (Santa Cruz Biotechnology, sc-13131), GAPDH (Santa Cruz Biotechnology, sc-47724), TIP60 (Santa Cruz Biotechnology, sc-166323), HUWE1 (Bethyl, A300-486A), Vinculin (Santa Cruz Biotechnology, sc-25336), 53BP1 (Bethyl, A300-272A), CTIP (Santa Cruz Biotechnology, sc-271339), LIG1 (Bethyl, A301-136A), PARP1 (Cell Signaling, 8542), PCNA (Cell Signaling, 2586), Cas9 (BioLegend 844302).

### Drug Sensitivity Assays

Olaparib was obtained from Selleck Chemicals and cisplatin was obtained from Biovision. To test cellular viability after olaparib treatment, cells were seeded in 96-well plates at a density of 2000 cells per well, treated with the indicated doses of olaparib for 3 days, and assessed using CellTiter-Glo reagent (Promega, G7572) per manufacturer’s instructions. Clonogenic survival assays were performed by seeding cells in 6-well plates; after 72 hours (olaparib) or 24 hours (cisplatin) of treatment at the indicated concentrations, media was changed and colonies were allowed to form for 2 weeks. Cells were fixed with a solution of 10% methanol + 10% acetic acid and stained using crystal violet (2% solution, Aqua Solutions). For apoptosis assays, cells were treated with olaparib (5 μM) for 3 days, prepared for flow cytometry using the FITC Annexin V kit (Biolegend, 640906) and quantified using a BD FACSCanto 10 flow cytometer.

### Functional assays

The chromatin immunoprecipitation (ChIP) experiments to investigate 53BP1 binding near a double-strand break site were performed as previously described with minor modifications.^24^ The U2OS-DSB reporter cell line, with an integrated reporter transgene and inducible expression of mCherry-LacI-FokI nuclease, were pretreated with the indicated siRNA for 48 hours, then treated with 4-hydroxytamoxifen (4-OHT, 1 μM) and shield-1 ligand (1 μM) to induce nuclease expression and subsequent double-strand break formation. Five hours after induction, cells were harvested and processed using the SimpleChIP Enzymatic Chromatin Immunoprecipitation Kit (Cell Signaling, 9003) according to the manufacturer’s instructions. The antibody used for immunoprecipitation was Anti- 53BP1 (Bethyl, A300-272A). Immunoprecipitated DNA was subjected to real-time qPCR with PerfeCTa SYBR Green SuperMix (Quanta), using a CFX Connect Real-Time Cycler (BioRad). Primers used for the DSB-reporter locus were:

for: GGAAGATGTCCCTTGTATCACCAT;
rev: TGGTTGTCAACAGAGTAGAAAGTGAA.

Detection of RPA-positive cells was performed as previously described, using a BD FACSCanto 10 flow cytometer.^29,30^ Data were analyzed using FlowJo software (BD). The neutral comet assay was performed per manufacturer’s instructions using the Comet Assay Kit (Trevigen, 4250-050) and olive tail moment was analyzed using CometScore. The DR-GFP homologous recombination assay was performed as previously described.^43^ DNA Fiber combing assays were performed as previously described.^16^

### Statistical analyses

For the neutral comet assays, the Mann-Whitney statistical test was performed. For all other assays, the statistical analysis performed was the t-test (two-tailed, unequal variance unless indicated). Statistical significance is indicated for each graph (ns = not significant, for P > 0.05; * for P ≤ 0.05; ** for P ≤ 0.01; *** for P ≤ 0.001; **** for P ≤ 0.0001).

### Genomic and transcriptomic analysis

Genomic and transcriptomic data for chemo-resistant, HGSC was obtained from the Australian Ovarian Cancer Study cohort (OV-AU), which is part of the International Cancer Genome Consortium (ICGC). Somatic structural variants (duplications, deletions, inversions, intra- and inter-chromosomal translocations) were identified using qSV^48^ from the whole-genome sequencing data, and length of homology was reported for the structural variation junctions. The structural variations were classified as non-homologous end-joining (NHEJ)-mediated if there was ≤3bp homologous sequences. There were 48–2,431 (median: 292) structural variants per sample in the ovarian cohort. RNA-sequencing-based expression data were also available for these tumors from the ICGC data portal (https://dcc.icgc.org/). We grouped the samples into four categories based on combinatorial high (above median) and low (below median) expression of BRCA2 and TIP60; this enabled us to analyze samples with BRCA deficiency due to classic BRCA2 inactivation or loss as well as those potentially mediated by other mechanisms. Survival analysis was performed based on the TCGA Ovarian cancer samples^49^ with and without BRCA2 mutations using ‘survival’ R package. Statistical analysis was performed using R.

## Acknowledgements

We would like to thank Jeremy Stark, Roger Greenberg, Robert Brosh, Mariano Russo, Jacob Hornick, Yunsung Kim, and Michael O’Connor for materials and advice; and the following Penn State College of Medicine core facilities: Flow Cytometry, Genomic Analyses, and Imaging. Experimental design schemes and summary models were created with Biorender.com. This work was supported by: NIH R01ES026184 and the St. Baldrick’s Foundation (to GLM), and NIH R01GM129066 (to SD).

