## Supplemental material for "Identification of regulators of poly-ADP-ribose polymerase (PARP) inhibitor response through complementary CRISPR knockout and activation screens"

**Legends to Supplemental Tables and Figures**

**Supplemental Table S1. Results of the CRISPR screens presented in Figure 1.** All genes targeted by the respective libraries are ranked according to P-value. Each gene was targeted by multiple sgRNAs within the libraries. The fold change in representation between the olaparib- and DMSO- treated conditions were calculated for each individual sgRNA, and these values were used to calculate a P-value for each gene using RSA analysis.

**Supplemental Figure S1. Gene ontology analysis of the top candidates from the CRISPR knockout screen for olaparib sensitivity in wildtype cells.** Top hits with LogP less than or equal to -2 were entered into NIH DAVID and pathways were analyzed for Gene Ontology Biological Processes (GO_BP) terms. GO_BP terms with a logP of -5 or lower are shown.

**Supplemental Figure S2. Gene ontology analysis of the top candidates from the CRISPR knockout screen for olaparib resistance in BRCA2$^{KO}$ cells.** Top hits with LogP less than or equal to -2 were entered into NIH DAVID and pathways were analyzed for GO_BP terms. GO_BP terms with a logP of -2.5 or lower are shown.

**Supplemental Figure S3. Gene ontology analysis of the top candidates from the CRISPR activation screen for olaparib resistance in BRCA2$^{KO}$ cells. (A)** Western blot confirming expression of dCas9 in HeLa BRCA2$^{KO}$ cells. **(B)** Hits with LogP less than or equal to -2 were entered into NIH DAVID and pathways were analyzed for GO_BP terms. GO_BP terms with a logP of -1.5 or lower are shown.

**Supplemental Figure S4. Knockdown of TIP60 or HUWE1 with either of two siRNA oligonucleotides causes PARPi resistance in HeLa BRCA2-knockout cells**. **(A)** Western blot showing that TIP60 is efficiently depleted with siRNA oligonucleotides employed. **(B)** Depletion of TIP60 by two siRNA oligonucleotides rescues PARPi sensitivity in HeLa BRCA2-knockout cells in cellular viability assays. **(C)** HUWE1 protein is depleted using two siRNA oligonucleotides as shown by western blot. **(D)** Depletion of HUWE1 by two siRNA oligonucleotides rescues PARPi sensitivity in HeLa BRCA2-knockout cells in cellular viability assays. **(E, F)** Olaparib-induced apoptosis is rescued by TIP60 **(E)** or HUWE1 **(F)** depletion in BRCA2-knockout cells. Asterisks indicate statistical significance compared to the olaparib-treated siControl condition. All data are shown as averages of three experiments with standard deviations as error bars.

**Supplemental Figure S5. TIP60 depletion does not affect other proposed mechanisms of PARPi cytoxicity. (A)** A chromatin fractionation experiment shows that while MMS (0.01%) and olaparib (1 μM) co-treatment for 3 hours induces trapping of PARP1 on the chromatin, this effect is not rescued by TIP60 depletion. **(B)** DNA fiber assay performed in HeLa BRCA2 knockout cells demonstrates that while olaparib treatment increases replication fork speed, this phenotype is not affected by TIP60 depletion. Olaparib (10μM) was added for 24 hours prior to incubation with the thymidine analogs. Horizontal bars reflect the means of replication fork speeds calculated from measurements of CldU tracts. At least 65 fibers were quantified from one experiment. Asterisks indicate statistical significance. **(C)** TIP60 depletion does not rescue the homologous recombination defect caused by BRCA2 depletion as shown using a DR-GFP assay. The averages of three experiments are shown, with standard deviations as error bars. Statistical significance is indicated by asterisks.

**Supplemental Figure S6. Functional impact of TIP60, LIG1, or CTIP depletion in BRCA2-knockout cells. (A)** Western blot showing that 53BP1 is depleted by the siRNA oligonucleotides employed, and that TIP60 depletion is equivalent when depleted alone or in combination with 53BP1. **(B)** Quantitative PCR assay demonstrating the efficacy of the siRNA oligonucleotide targeting REV7. Data shown is the average of three experiments, normalized to the siControl condition, with standard deviations shown as error bars. Statistical significance is indicated by asterisks. **(C)** Flow cytometry dot plot of DAPI versus RPA staining intensity illustrates the gating (inset box) used to produce Figure 5B. Cells in Quadrant 2 (Q2) were considered RPA-positive, and cells in Quadrant 3 (Q3) RPA-negative; this cutoff was determined based on the “No treatment” condition and was kept constant across all samples. Approximately 20,000 events per sample from a single experiment are shown, and similar results were obtained across 3 experiments. **(D)** The efficacy of the siRNA oligonucleotides targeting CTIP and LIG1 are shown using western blot. **(E)** Cellular viability assays demonstrate that depletion of CTIP or LIG1 is not sufficient to cause PARPi resistance in BRCA2-knockout HeLa cells. The averages of three experiments are shown, with standard deviations as error bars.

### Supplemental Figure S1

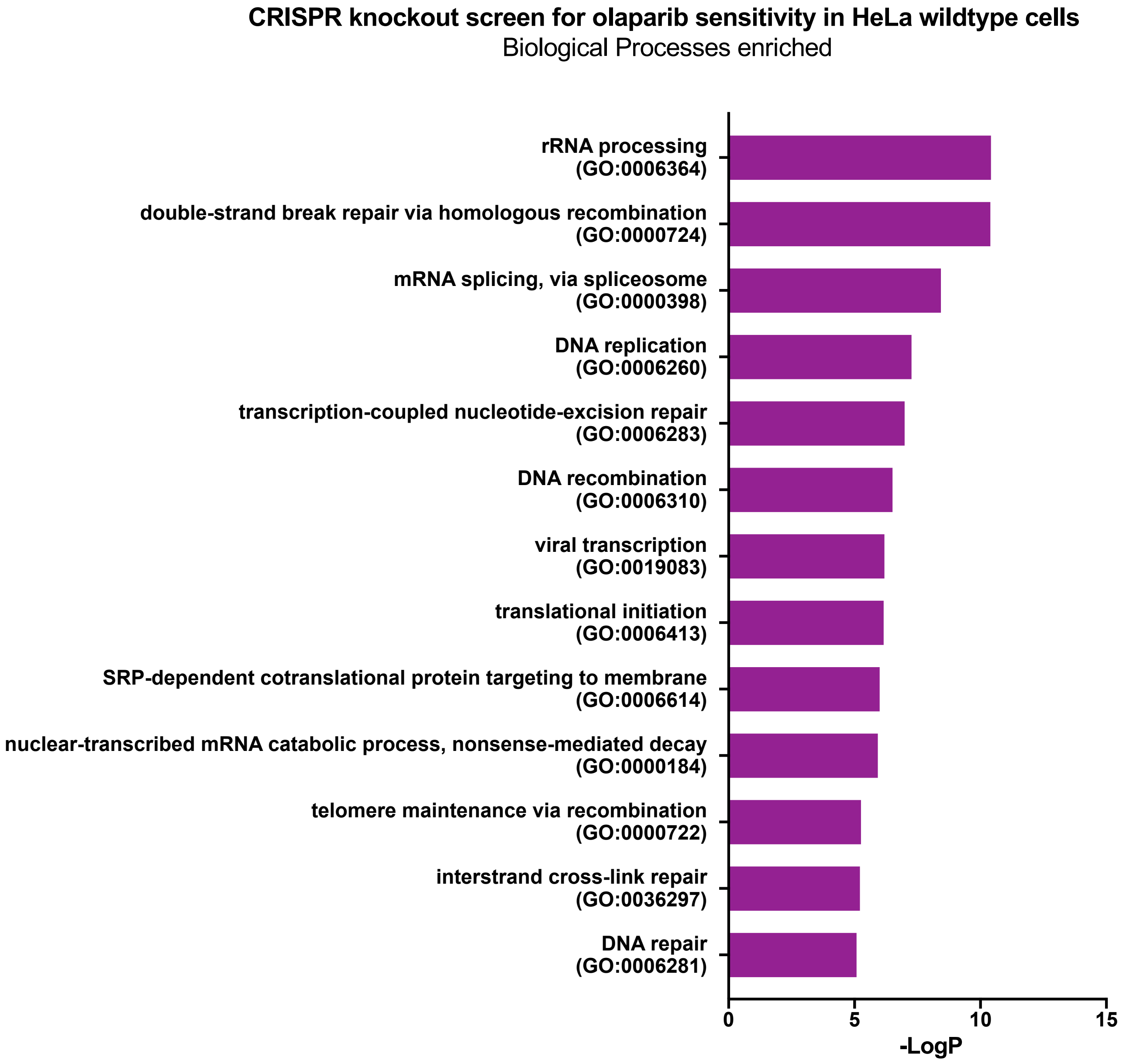

### Supplemental Figure S2

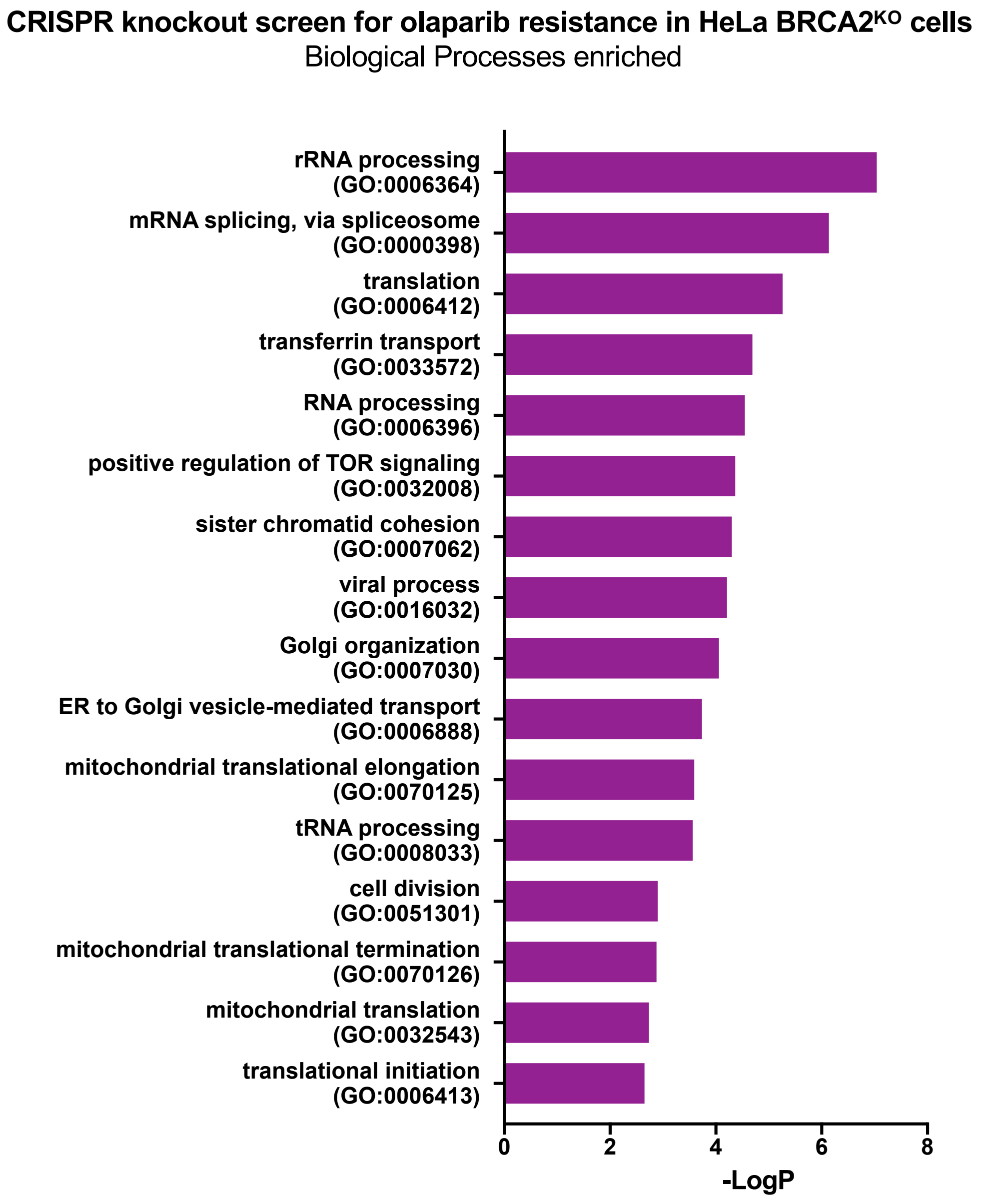

### Supplemental Figure S3

**A**

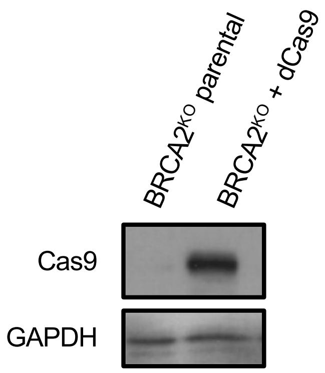


**B**

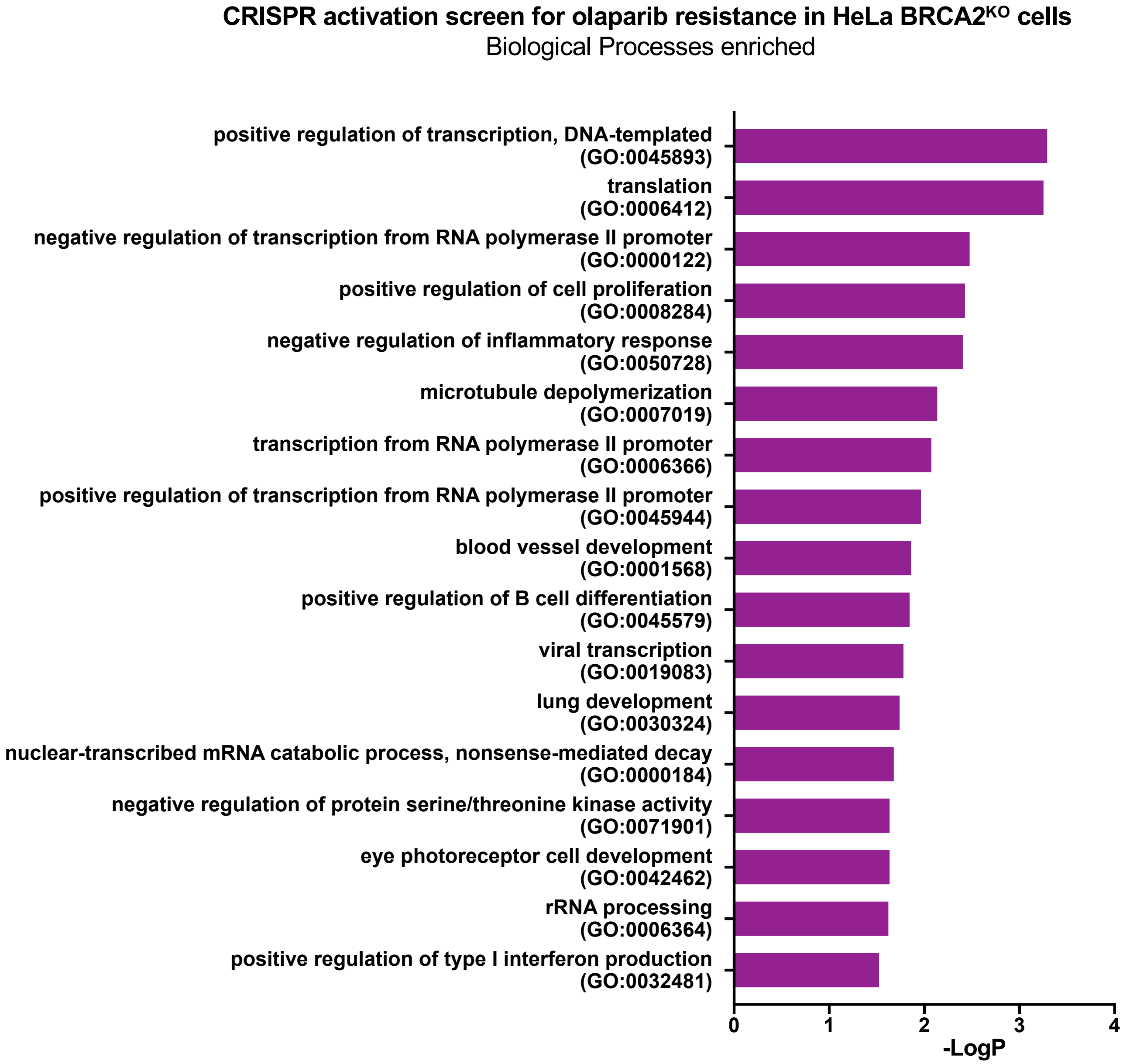

### Supplemental Figure S4

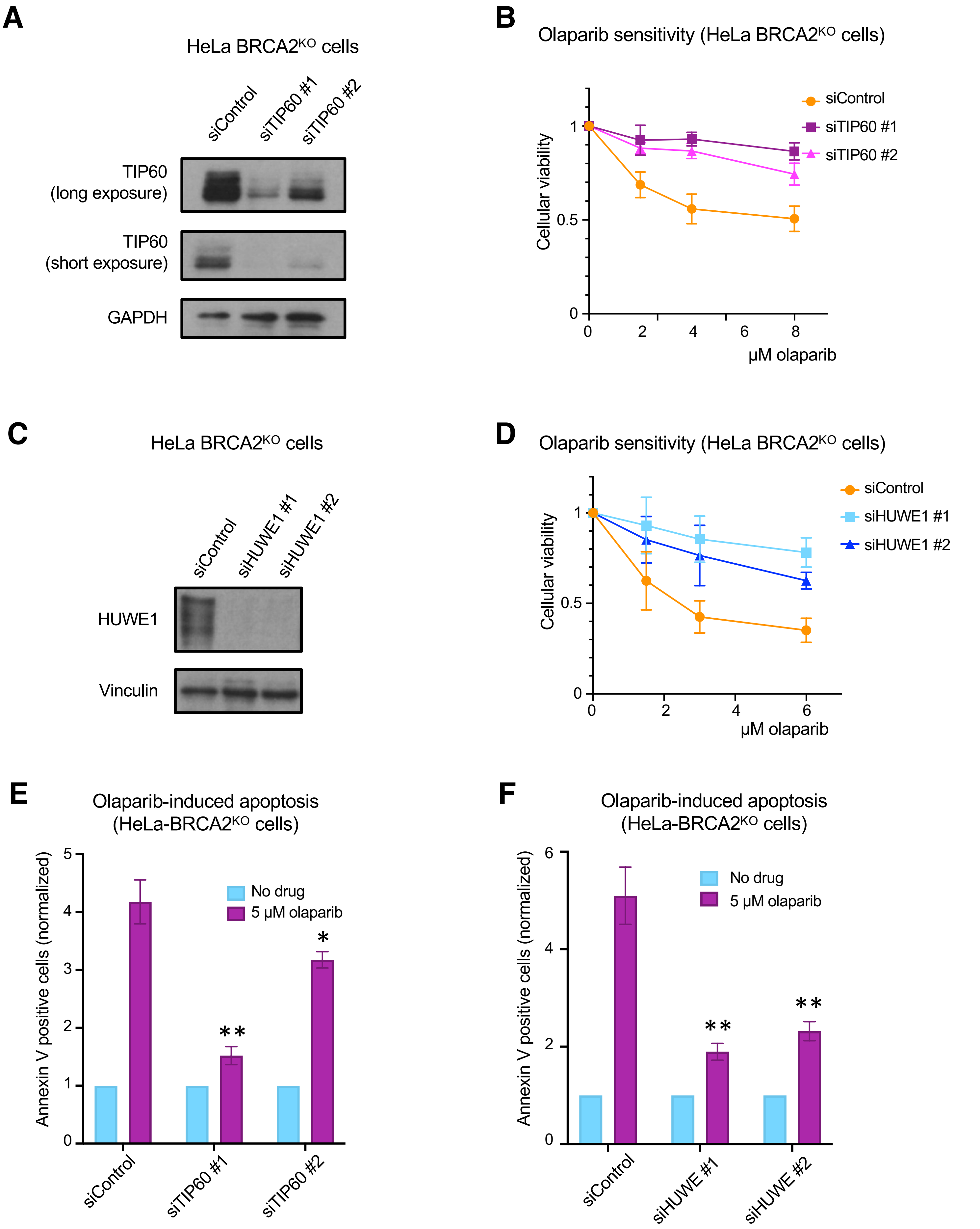

### Supplemental Figure S5

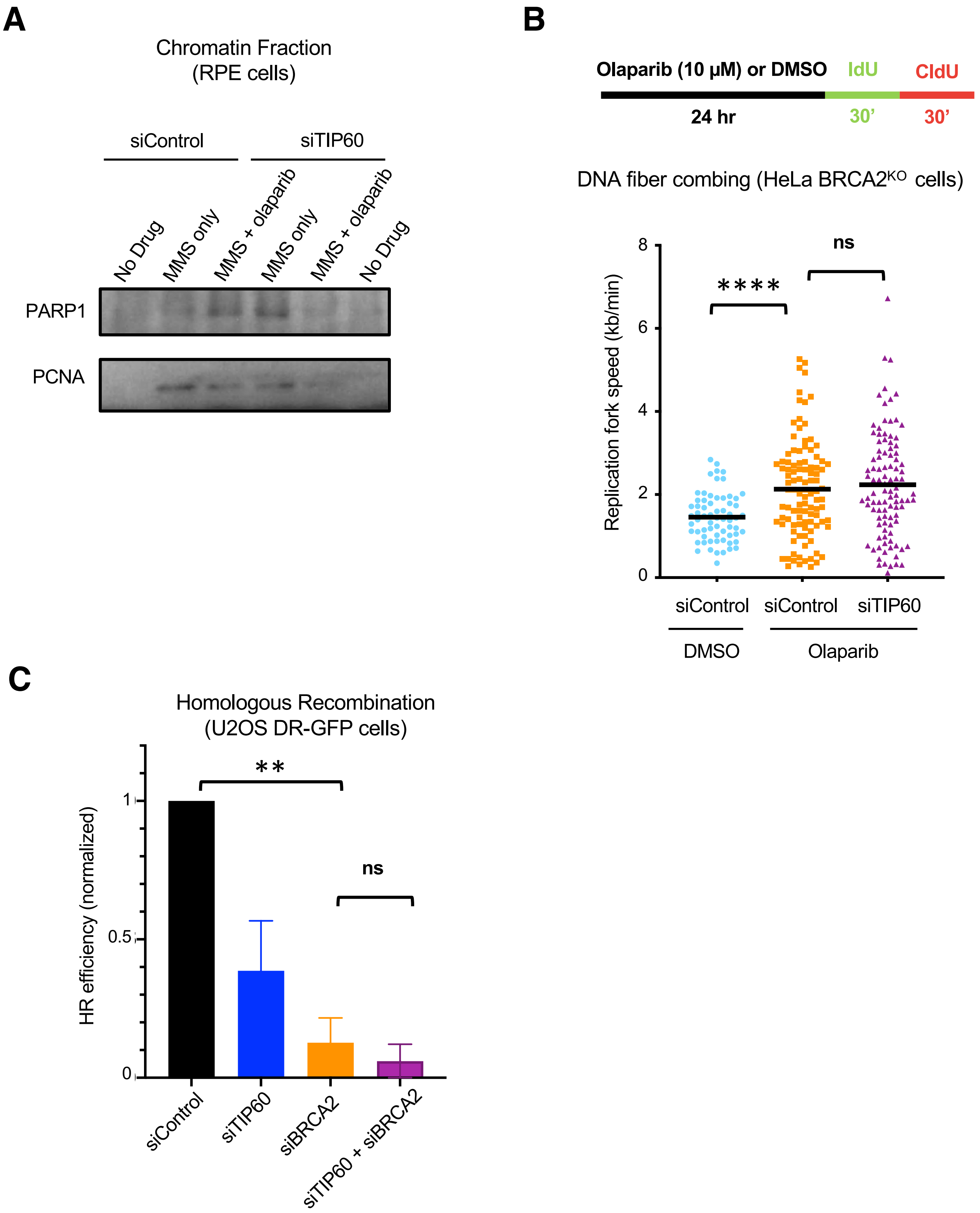

### Supplemental Figure S6

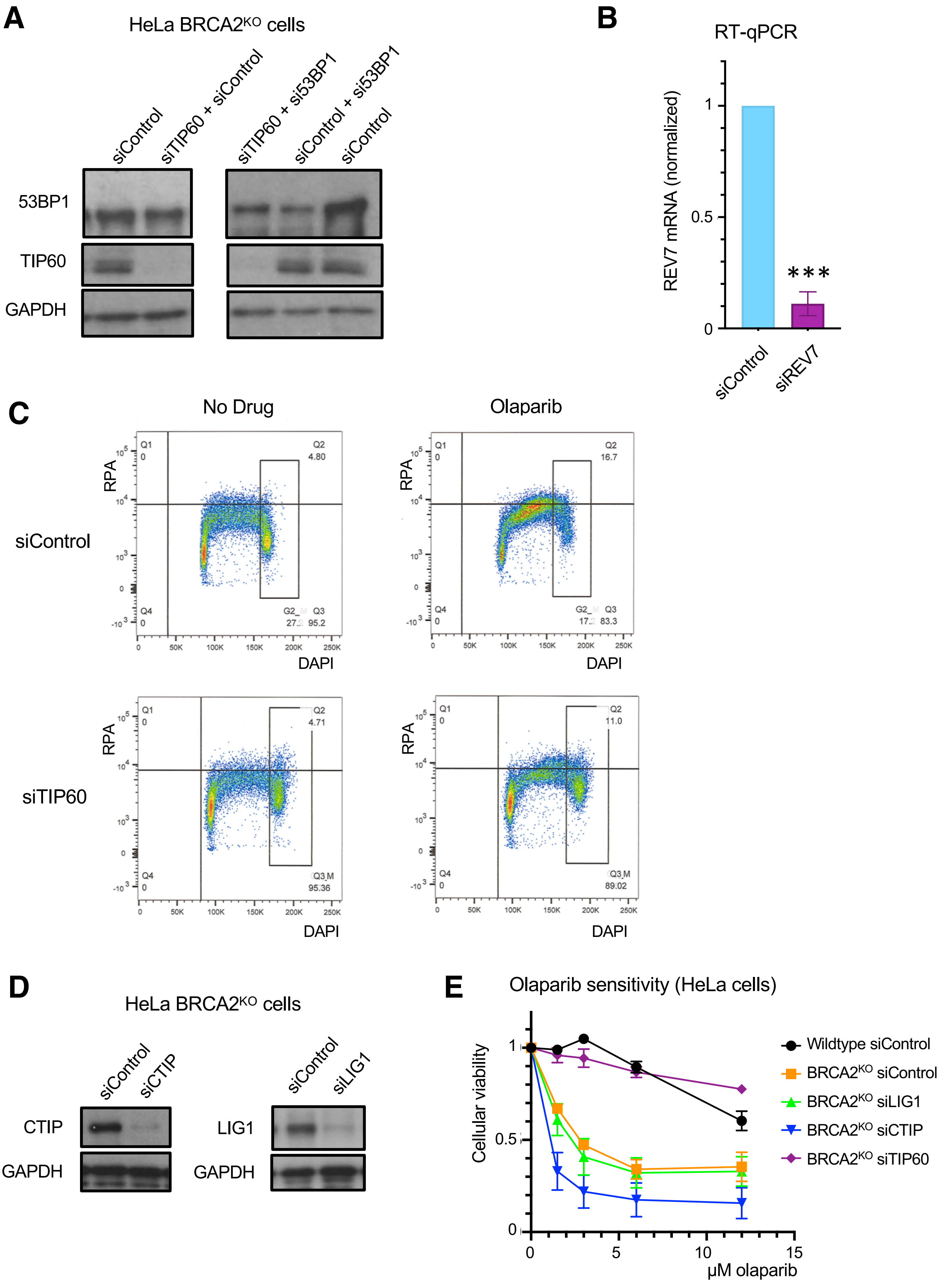
